# Enzyme Co-Scientist: Harnessing Large Language Models for Enzyme Kinetic Data Extraction from Literature

**DOI:** 10.1101/2025.03.03.641178

**Authors:** Jinling Jiang, Jie Hu, Siwei Xie, Menghao Guo, Yuhang Dong, Shuai Fu, Xianyue Jiang, Zhenlei Yue, Junchao Shi, Xiaoyu Zhang, Minghui Song, Guangyong Chen, Hua Lu, Xindong Wu, Pei Guo, Da Han, Zeyi Sun, Jiezhong Qiu

## Abstract

The extraction of molecular annotations from scientific literature is critical for advancing data-driven research. However, traditional methods, which primarily rely on human curation, are labor-intensive and error-prone. Here, we present an LLM-based agentic workflow that enables automatic and efficient data extraction from literature with high accuracy. As a demonstration, our workflow successfully delivers a dataset containing over 91,000 enzyme kinetics entries from around 3,500 papers. It achieves an average F1 score above 0.9 on expert-annotated subsets of protein enzymes and can be extended to the ribozyme domain in fewer than 3 days at less than $90. This method opens up new avenues for accelerating the pace of scientific research.

## Main Text

In scientific research, acquiring a vast amount of annotations or “labels” for functional molecules enables the identification of subtle patterns and the development of predictive models. Increasing availability of human-curated scientific datasets facilitates the integration of interdisciplinary insights, driving innovation and accelerating discovery in multiple fields including chemistry[1], biology[2-5], and material science[6]. By the year 2025, the Nucleic Acids Research (NAR) Database issue has collected up to 2236 databases[7], among which many are constructed through human-curated literature mining such as ChEMBL[8] and BRENDA[9]. Robust data-driven research will foster more efficient scientific progress, but traditional methods by human annotation and classical natural language processing tools are labor-intensive, time-consuming, and error-prone[10]. Therefore, there is an urgent need for paradigm-shifting methods capable of rapidly and accurately extracting annotations from literature.

Recent advancements in Artificial General Intelligence (AGI), particularly Large Language Models (LLMs), have demonstrated superior performance in scientific literature mining such as named entity recognition, relation extraction, summarization, integration and even planning[5,11-16]. Most recently, OpenAI has unveiled ‘deep research’ that can conduct multi-step research on the internet for complex tasks to assist scientists to write literature reviews and even identify knowledge gaps[17], and Google has launched ‘AI co-scientist’ which is “*a virtual scientific collaborator to help scientists generate novel hypotheses and research proposals and to accelerate the clock speed of scientific and biomedical discoveries*”[18].

Nevertheless, extracting data—especially quantitative annotations—from literature that contains both tables and text remains challenging. This task requires comprehensive long-context capabilities, including long-context retrieval and reasoning, tabular data understanding, and basic math (such as scientific notation understanding). Therefore, mining quantitative data from literature naturally serves as a real-world, large-scale benchmark for evaluating the long-context capabilities of LLMs. This is particularly significant given that most existing long-context LLM benchmarks[19,20] rely on synthetic data or costly human annotations.

To address the aforementioned challenges, we have developed an agentic workflow for automated extraction, normalization, and standardization of scientific data from the literature (Fig. 1). The agentic workflow starts with collecting a full dataset of relevant literature in PDF format, which are then transformed to machine-readable text by optical character recognition (OCR). Then the text, together with an engineered prompt, is used to query LLMs to extract data of interest. The outputs of multiple LLMs are then considerably aggregated to provide comprehensive and high-confidence results.

**Fig. 1.**
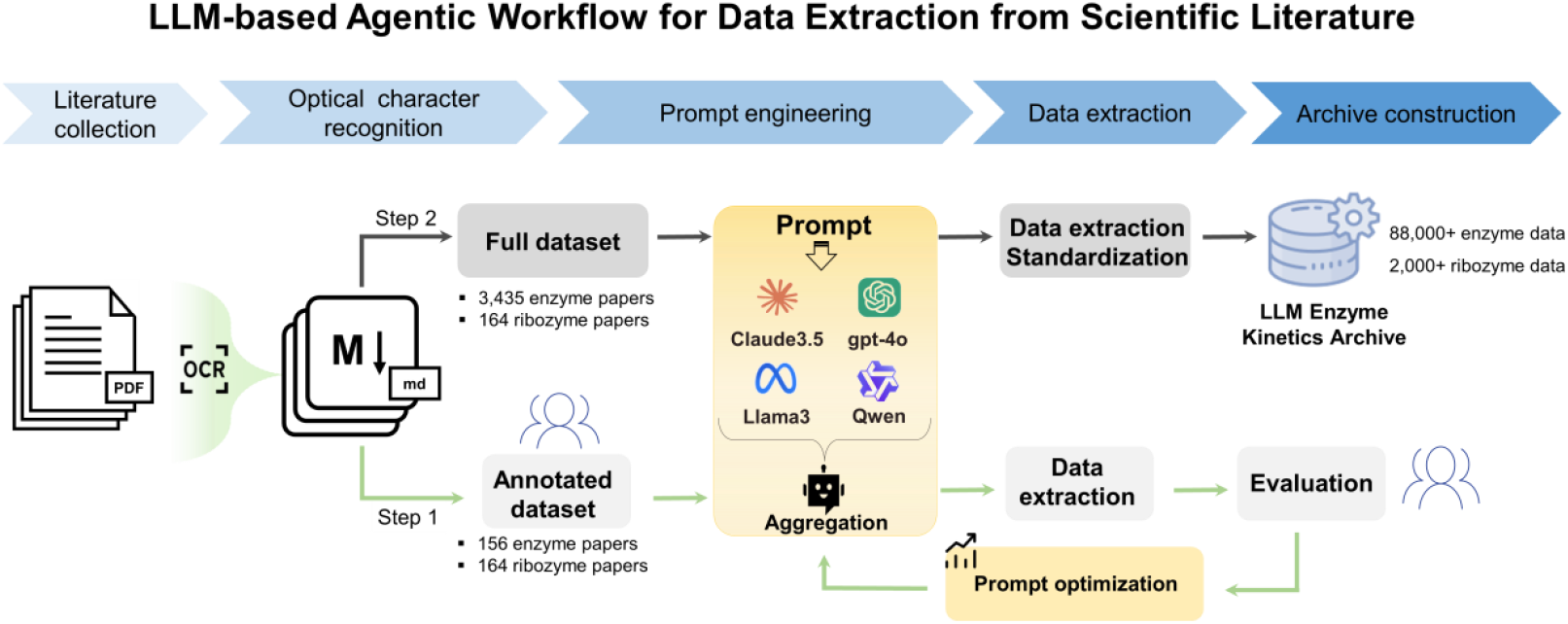
Schematic of our LLM-based agentic workflow for enzyme kinetic data extraction. To evaluate and further optimize our workflow, we manually annotate a small portion of the full dataset (termed as the annotated dataset) to thoroughly evaluate each individual component of the workflow, including the prompt, choice of LLMs, and aggregation strategy. First, the prompt is carefully engineered to include task background, field description, output format instruction, etc, which are essential for LLMs to process desired information with high accuracy. Second, numerous LLMs have been developed, each with its own strengths and weaknesses. Benchmarking different LLMs allows for a comprehensive evaluation of their capabilities across various dimensions, including long-context reasoning, information retrieval, coreference resolution, unit conversion, and table understanding. Finally, an LLM aggregator is designed to take the strengths of high-performance LLMs towards higher confidence results.

We first apply our workflow to extract kinetic data of protein enzymes, which is an important domain in the field of biochemistry with well-established databases such as BRENDA[9]. This allows us to thoroughly assess the performance of our workflow compared to existing human-curated databases. Following the workflow in Fig. 1, we extract three types of kinetic parameters, including K_m_, *k*_cat_ and *k*_cat_/K_m_, from 3,435 papers curated in BRENDA.

To rigorously evaluate extraction performance, we randomly select 156 papers and manually annotate them with a team of experts, resulting in an annotated dataset that contains 3,563 K_m_, 3,531 *k*_cat_, and 3,751 *k*_cat_/K_m_ entries. The extracted results are compared to the expert annotation to calculate metrics including paper-wise precision, recall, and F1-score (Methods, Supplementary Table 1). We evaluate four popular LLMs, including two closed-source ones (Claude3.5 and gpt-4o) and two open-source ones (Llama3 and Qwen), with our engineered prompt (Fig. 2a, d). Claude3.5 is identified as the top performer in terms of F1-score distribution (mean=0.90, median=0.99, 25%/75% percentile=0.90/1.00) and exhibits more consistent performance across different temperature settings (Supplementary Fig. 1). However, we observe that, despite their lower mean and median F1 scores, the other three LLMs demonstrate distinct strengths in resolving certain challenging cases encountered by Claude3.5 (Examples of these challenging cases are discussed in Supplementary Note 1). Inspired by ensemble learning, we develop an aggregation agent to leverage the strengths of individual LLMs. In particular, we prompt Claude3.5 with the paper text, along with the outputs from Claude3.5, gpt-4o, Llama3, and Qwen, and then instruct it to consolidate the insights from these LLMs (Fig. 2e). Encouragingly, the aggregation agent is indeed capable of integrating the advantages of multiple LLMs and achieves a comprehensively enhanced performance, especially in reducing bad cases (Fig. 2b). Finally, we investigate whether LLM could surpass human intelligence in literature mining by comparing results produced by LLMs with data curated in BRENDA (Methods). For the 156 papers in our annotated dataset, BRENDA achieves suboptimal F1 scores (Fig. 2c, mean=0.76, median=0.78, 25%/75% percentile=0.64/0.95) compared to any single LLM, or the advanced aggregation agent. We also validate the performance of our pipeline on the full dataset, comprising 3,435 papers, which shows that our pipeline maintains a quality comparable to that of the annotated dataset (Supplementary Note 2).

**Fig. 2.**
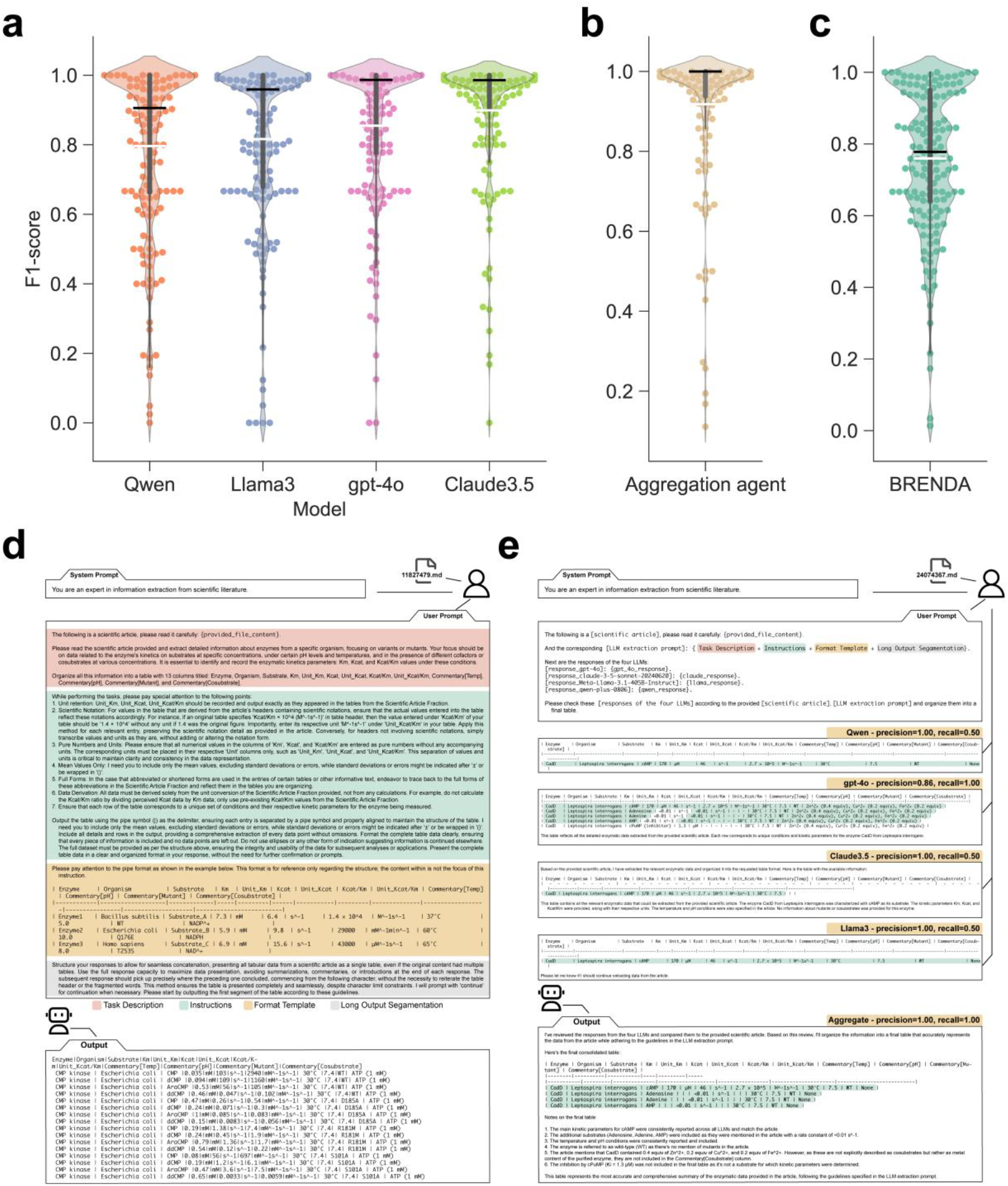
Protein Enzyme Kinetic Data Extraction. **a-c**, Violin plots show the distribution of paper-wise F1 scores of each LLM, sorted by mean F1 (**a**), the aggregation agent that integrates Claude3.5, gpt-4o, Llama3, and Qwen (**b**), and BRENDA (**c**) on the expert-annotated dataset (156 papers). Black bar: median. White bar: mean. 25% and 75% percentile and whiskers are also plotted. **d-e**, The engineered prompts for running single LLM (**d**) and aggregation agent (**e**).

As an important complement to enzymes, certain RNA molecules, known as ribozymes, possess catalytic activities and pivotal functions in gene regulation. However, to the best of our knowledge, there is no database service available for ribozyme kinetic data. Therefore, we next apply our workflow to the ribozyme research field where a brand-new database needs to be established from scratch. We collect a total of 164 papers from PubMed and bioRxiv by searching the keywords ‘ribozymes and (*k*_cat_ or K_m_ or *k*_cat_/K_m_ or *k*_obs_ or *k*_cleave_)’, which can be approximately considered as the full dataset of ribozyme kinetic literature. We apply our pipeline to extract five kinetic parameters, including *k*_cat_, K_m_, *k*_cat_/K_m_, *k*_obs_, and *k*_cleave_, and compare the extracted results with expert annotation (Supplementary Table 2). Again, Claude3.5 shows an exceptional performance with the highest mean F1 score of 0.75 (Fig. 3a). An aggregation agent that integrates these four LLMs does not perform better than Claude3.5, which is probably lagged by the poorer performances of the other three LLMs, especially gpt-4o and Qwen with mean F1 scores below 0.6. However, integrating Claude3.5 and Llama3 can achieve superior performance (Fig. 3b), which suggests the need for a smarter aggregation agent that should automatically vote out poorly performed LLMs during ensemble. Remarkably, our pipeline is economical and time-saving, i.e., it takes less than 90$ (Fig. 3c) to deliver the final extraction results within 3 days (2 days for literature collection and 1 day for running OCR software and LLMs). This demonstrates the efficiency of our workflow in acquiring knowledge for a specific research field with significantly improved data quality and reduced labor cost.

**Fig. 3.**
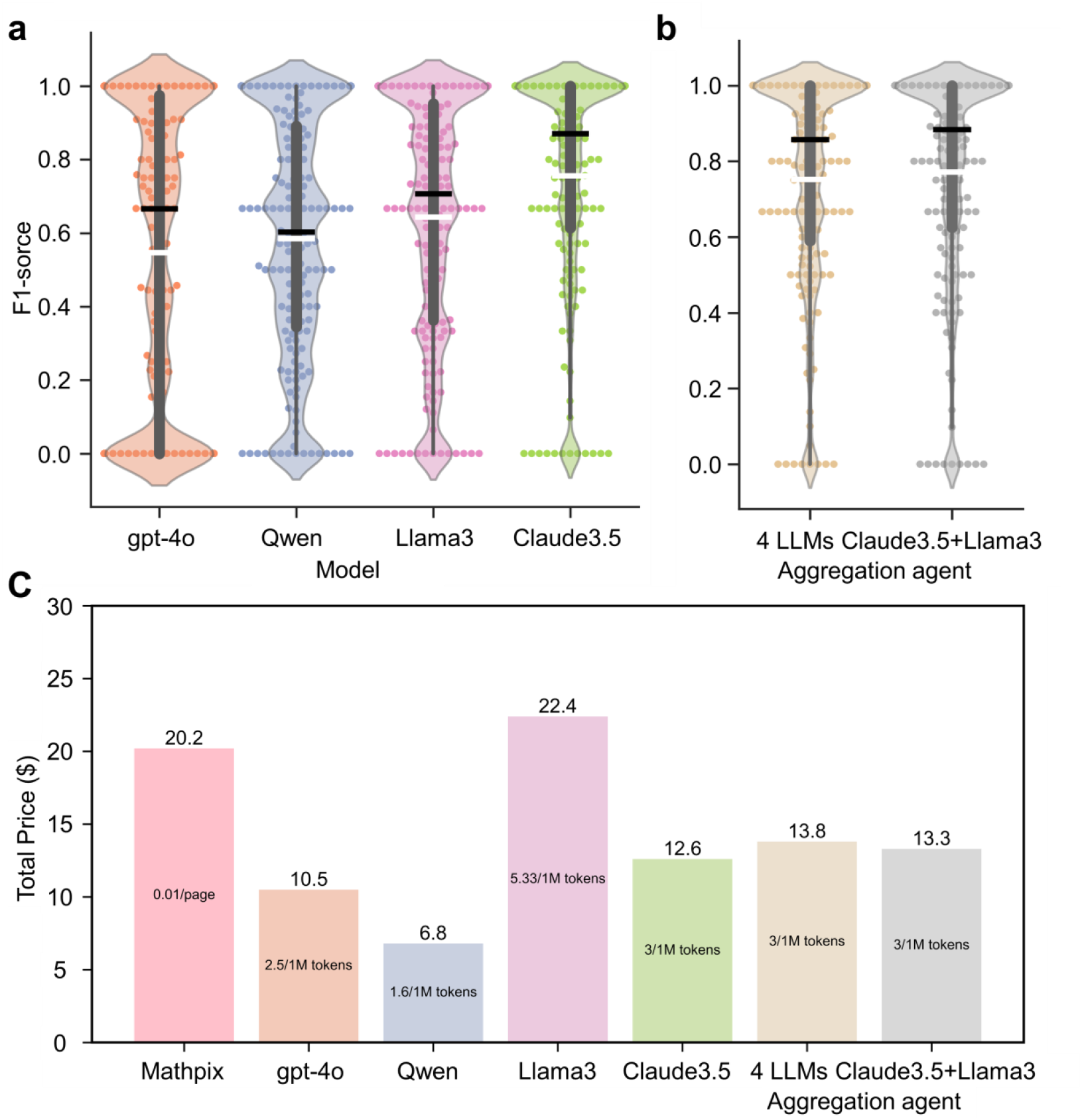
Ribozyme kinetic data extraction. **a**, Violin plots show the distribution of paper-wise F1 scores of each LLM against our expert-annotated dataset sorted by mean F1 (164 ribozyme papers). **b**, The performance of aggregation agent that integrates four LLMs including Claude3.5, gpt-4o, Llama3, and Qwen (left), and two LLMs including Claude3.5 and Llama3 (right). Black bar: median. White bar: mean. 25% and 75% percentile and whiskers are also plotted. **c**, The per-step cost of our pipeline when processing the 164 ribozyme papers.

To the end, we release a new archive^1^ that offers a structured and accessible collection of enzyme kinetic data, including 88,770 entries from 3,435 protein enzyme papers and 2,420 entries from 164 ribozyme papers (Supplementary Fig. 2). The archive contains kinetic parameters with their corresponding enzyme names, mutants, organisms, substrates, and experimental conditions, offering a valuable resource to the field of biochemistry. In addition, this archive serves as a real-world large-scale long-context LLM benchmark, with an average token count of 25k (385 papers containing over 32k tokens), allowing us to assess LLMs’ capabilities in understanding long scientific literature.

In sum, we have developed an LLM-based agentic workflow for automatic data extraction from scientific literature, providing a streamlined pipeline that operates without strong assumptions or prior knowledge for the domain of interest. By demonstrating the high accuracy and efficiency of our workflow in the domains of protein enzyme and ribozyme, we anticipate it can also be robustly adapted to other research fields. This work validates and benchmarks the superiority of LLMs in literature mining compared to human efforts. In the future, we aim to refine the enzyme archive to facilitate new discoveries and further extend our workflow to other fields towards a versatile paradigm and a real LLM benchmark in scientific research.

## Methods

### Literature Collection

The papers about protein enzyme are collected from BRENDA whose core information is manually extracted from original scientific literature from PubMed and Scopus. We get a collection of PubMed Identifier (PMID) from BRENDA database through the BRENDA SOAP (Simple Object Access Protocol) services. The number of PMIDs we collect from BRENDA is more than 90,000 among which 4,925 papers have recorded the experimental results of enzyme kinetics parameters. We finally download 3,435 papers in PDF format by PMIDs as we do not have access to the remaining 1,490 ones. For these papers, we implement a crawler program to download their corresponding enzyme kinetic data from BRENDA.

As for the ribozyme, the 164 papers are collected from PubMed and bioRxiv by searching the keywords ‘ribozymes and (*k*_cat_ or K_m_ or *k*_cat_/K_m_ or *k*_obs_ or *k*_cleave_)’.

### Construction Expert-Annotated Datasets

We engage a team of 10 highly qualified experts to meticulously annotate, with 6 of them holding PhD degrees in relevant fields and the remaining 4 pursuing their PhDs in related disciplines. Through the annotation process, we ensure precision at every step, employing rigorous validation and repeated confirmations. We designate this as our gold standard for benchmarking, which we believe is exceptionally close to the true values. The final annotated dataset is structured as a wide table with aligned fields, including enzyme, mutant, organism, substrate, experimental conditions (such as temperatures and pHs), and kinetic data fields.

### OCR Pre-Processing

OCR is performed to convert PDF format to text format while preserving mathematical content and complex tables. We compare two OCR approaches: Mathpix and PyMuPDF, where we use Claude3.5 as the default LLM to extract protein enzyme data from the annotated datasets. As shown in Supplementary Fig. 3, Mathpix achieves a similar median F1 score compared to PyMuPDF (0.99 vs 1.00) but demonstrated a slightly better mean F1 score (0.90 vs 0.87). Therefore, we select Mathpix as the default OCR software in our pipeline.

### Prompt Engineering

The procedure of prompt engineering contains several key steps:

1. Evaluate the prompt using all samples in the annotated dataset.
2. Manually analyze problematic cases from samples with low precision or recall, identifying common errors.
3. For each identified error, we manually update the instruction part of the prompt.

The optimized prompt comprises three key components: task description, instructions, and format reference. The task description specifies the fields to be extracted. The instructions integrate essential reminders from the prompt engineering process. The format reference provides a template for the expected output. We selected Markdown as the output format because it avoids issues with separators (e.g., commas in compound names) and enables direct table visualization during prompt engineering, a feature lacking in other formats like JSON. The final prompt is listed in Supplementary Note 3. The prompt for the aggregation agent is listed in Supplementary Note 4.

### Post-Processing

The post-processing program organizes and cleans the outputs of LLMs, converting them into a standardized data frame format. This transformation enables downstream numerical comparison and evaluation. The program utilizes the Pandas library to structure the extracted data, primarily focusing on the outputs that present a table with 13 columns delineated by pipe delimiters. It removes any non-tabular descriptive content, aligns column headers, and implements necessary data-cleaning procedures while maintaining the integrity of the dataset. The cleaning procedures consists of several critical tasks:

1. Space cleaning. It removes undesired non-breaking spaces, fills empty entries with NA (Not Applicable) to avoid leaving blank spaces, and eliminates any rows where all entries are marked as NA.
2. Invalid field cleaning. Values under certain columns carrying units corresponding to other parameters will be marked as NA.
3. Numerical cleaning. It employs regular expressions to identify various representations of entries with scientific notations and/or error metrics.
4. Unit conversion. It involves recognizing the extracted units and standardizing the values of K_m_, *k*_cat_, and *k*_cat_/K_m_ to mM, s^-1, and s^-1mM^-1, respectively, by applying specified conversion factors.

This program transforms raw data extracted from LLM outputs into a clean and standardized format, ensuring data consistency and facilitating comparability for subsequent evaluations and applications.

### Evaluation Metrics

The performance metrics (precision, recall, and F1) are computed based on the kinetic constants (i.e., *k*_cat_, K_m_, *k*_cat_/K_m_, *k*_obs_, and *k*_cleave_), including both their numerical values and corresponding units. We observe that it is highly unlikely for a paper to contain multiple data entries with identical kinetic constants, even when papers have dozens or hundreds of entries. In other words, these constants can function as “unique identifiers,” reducing the need for additional processing after extraction and thereby simplifying the automatic evaluation process. More formally, the paper-wise precision of each paper in the annotated dataset is calculated using the formula:

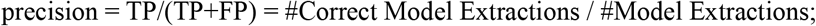

Similarly, the paper-wise recall is calculated as:

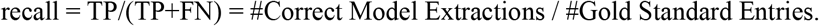

The F1 score is the harmonic mean of the precision and recall:

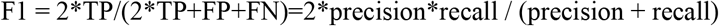

In some rare cases, the calculation of the above metrics can cause an error of division by zero. Take the computation of precision for example, this can happen if the LLM fails to extract any results, which makes the true positive (TP) as well as the false positive (FP) zero. For these special cases, we follow the common strategy:

- If the true positive (TP), false positive (FP), and false negative (FN) are all zeros, the precision, recall, and F1 are 1. This means that the LLM correctly knows “it doesn’t know” when there is indeed nothing to be extracted.
- If true positives (TP) is zero and at least one of the two others (i.e., FP and FN) is not, the precision, recall and F1 are all zeros.

Supplementary Tables 1 and 2 show the average metrics across all protein enzyme papers and ribozyme ones, respectively. We consider two ways to average --- macro and micro. The macro-averaged metrics (or macro metrics) are computed by taking the arithmetic mean (a.k.a. unweighted mean) of all the paper-wise metrics. For micro metrics, the counters (i.e., true positive, false positive, false negative) of all papers are summed up, which are then used to calculate a single micro precision/recall/F1 value.

However, the above metrics may not be directly applicable when comparing to data entries recorded in BRENDA, as BRENDA involves manual extraction where different levels of rounding are applied. For example, a K_m_ value of 0.835 might be rounded to 0.8 or 0.84 in BRENDA. Therefore, when comparing with BRENDA, we consider an extraction to be correct if rounding to any position between the 1st and 6th decimal place results in an exactly matched value.

### Details of LLMs Used

1. Version of LLMs: Claude-3.5-sonnet-20240620, Qwen-plus-0806, gpt-4o (September 23, 2024), Llama-3.1-405B.
2. Llama3 is locally deployed, while the other LLMs were used through online APIs.
3. Claude3.5 uses a temperature of 0.1. Other LLMs use a temperature of 0.3.
4. The maximum outputs of different LLMs vary, which is discussed in our paper: gpt-4o’s output capacity is 4096 tokens; Claude3.5’s output capacity is 8192 tokens; Qwen-Plus’s output capacity is 8000 tokens, and Llama3’s output capacity is 4096 tokens.
5. Due to local GPU resource limitations, Llama3 used a maximum input of 32k tokens.

## Data Availability

Our annotated dataset (156 protein enzyme papers and 164 ribozyme papers) is available at https://huggingface.co/datasets/jackkuo/LLM-Enzyme-Kinetics-Golden-Benchmark. Our released enzyme archive (88,770 entries from 3,435 protein enzyme papers and 2,420 entries from 164 ribozyme papers) is available at https://huggingface.co/datasets/jackkuo/LLM-Enzyme-Kinetics-Archive-LLENKA.

## Code Availability

All codes used in this study have been deposited on GitHub: https://github.com/JackKuo666/LLM-BioDataExtractor

## Acknowledgments

This research is supported by the National Natural Science Foundation of China (62120106008, 62306290), the “Pioneer” and “Leading Goose” R&D Program of Zhejiang (2024SSYS0007). We would like to express our gratitude to the Qwen Team at Alibaba for providing access to the free Qwen API.

## Footnotes

1 https://huggingface.co/datasets/jackkuo/LLM-Enzyme-Kinetics-Archive-LLENKA

