## Supplementary Information for "Enzyme Co-Scientist: Harnessing Large Language Models for Enzyme Kinetic Data Extraction from Literature"

[1]Zhejiang Lab, Hangzhou, Zhejiang, China.

[2]Hangzhou Institute of Medicine Chinese Academy of Sciences, Hangzhou, Zhejiang, China.

[3]Department of Computer Science, Aalborg University, Denmark.

[4]Key Laboratory of Knowledge Engineering with Big Data (The Ministry of Education of China), Hefei University of Technology, Hefei, Anhui, China.

[#]Authors who contribute equally.

**Supplementary Table 1.** Overall performance of various models examined on the annotated dataset of 156 protein enzyme papers.

| | | | Llama 3 | Claude3.5 | gpt-4o | Qwen | BRENDA | Aggre |
|---|---|---|---|---|---|---|---|---|
| Overall (10845) | Micro | Recall | 0.72 | **0.92** | 0.80 | 0.72 | 0.77 | 0.90 |
| | | Precision | 0.90 | 0.88 | 0.89 | 0.90 | 0.75 | **0.93** |
| | | F1 | 0.80 | 0.90 | 0.84 | 0.80 | 0.76 | **0.91** |
| | Macro | Recall | 0.80 | 0.93 | 0.85 | 0.77 | 0.80 | **0.94** |
| | | Precision | 0.89 | 0.90 | 0.90 | 0.90 | 0.77 | **0.93** |
| | | F1 | 0.82 | 0.90 | 0.85 | 0.80 | 0.76 | **0.92** |
| $K_m$ (3563) | Micro | Recall | 0.74 | **0.93** | 0.82 | 0.75 | 0.82 | 0.91 |
| | | Precision | 0.92 | 0.86 | 0.89 | 0.91 | 0.82 | **0.93** |
| | | F1 | 0.82 | 0.90 | 0.85 | 0.80 | 0.76 | **0.92** |
| | Macro | Recall | 0.83 | **0.94** | 0.88 | 0.79 | 0.85 | **0.94** |
| | | Precision | 0.90 | 0.90 | 0.90 | 0.89 | 0.83 | **0.93** |
| | | F1 | 0.84 | 0.91 | 0.87 | 0.81 | 0.82 | **0.92** |
| $\boldsymbol{k_{cat}}$ (3531) | Micro | Recall | 0.73 | **0.94** | 0.82 | 0.75 | 0.78 | 0.91 |
| | | Precision | 0.91 | 0.89 | 0.91 | 0.91 | 0.83 | **0.93** |
| | | F1 | 0.81 | 0.91 | 0.86 | 0.82 | 0.80 | **0.92** |
| | Macro | Recall | 0.81 | 0.92 | 0.86 | 0.78 | 0.81 | **0.94** |
| | | Precision | 0.89 | 0.90 | 0.90 | 0.88 | 0.85 | **0.93** |
| | | F1 | 0.83 | 0.90 | 0.86 | 0.80 | 0.80 | **0.93** |
| $\boldsymbol{k_{cat}/K_m}$ (3751) | Micro | Recall | 0.69 | **0.89** | 0.77 | 0.66 | 0.71 | 0.88 |
| | | Precision | 0.87 | 0.89 | 0.87 | 0.87 | 0.63 | **0.92** |
| | | F1 | 0.77 | 0.89 | 0.81 | 0.75 | 0.67 | **0.90** |
| | Macro | Recall | 0.79 | **0.92** | 0.84 | 0.74 | 0.75 | **0.92** |
| | | Precision | 0.86 | 0.89 | 0.87 | 0.83 | 0.68 | **0.90** |
| | | F1 | 0.80 | 0.89 | 0.84 | 0.76 | 0.69 | **0.90** |

**Supplementary Table 2.** Overall performance of various models examined on the annotated dataset of 164 ribozyme papers.

| | | | Llama 3 | Claude3.5 | gpt-4o | Qwen | Aggre - 4 LLMs | Aggre - Claude3.5+Llama3 |
|---|---|---|---|---|---|---|---|---|
| Overall (2420) | Micro | Recall | 0.58 | 0.76 | 0.51 | 0.59 | 0.76 | **0.78** |
| | | Precision | 0.66 | **0.79** | 0.77 | 0.49 | 0.76 | 0.78 |
| | | F1 | 0.62 | **0.78** | 0.61 | 0.54 | 0.76 | **0.78** |
| | Macro | Recall | 0.67 | 0.80 | 0.53 | 0.67 | 0.81 | **0.83** |
| | | Precision | 0.69 | 0.75 | 0.62 | 0.60 | 0.75 | **0.76** |
| | | F1 | 0.64 | 0.76 | 0.55 | 0.59 | 0.75 | **0.77** |
| $\boldsymbol{k_{obs}}$ (1100) | Micro | Recall | 0.53 | 0.77 | 0.38 | 0.56 | 0.74 | **0.78** |
| | | Precision | 0.70 | 0.75 | **0.78** | 0.53 | 0.74 | 0.76 |
| | | F1 | 0.60 | 0.76 | 0.51 | 0.54 | 0.74 | **0.77** |
| | Macro | Recall | 0.67 | 0.79 | 0.58 | 0.67 | **0.80** | **0.80** |
| | | Precision | 0.72 | 0.78 | 0.66 | 0.68 | **0.80** | 0.79 |
| | | F1 | 0.67 | 0.76 | 0.60 | 0.63 | **0.78** | **0.78** |
| $K_m$ (404) | Micro | Recall | 0.74 | 0.91 | 0.71 | 0.82 | **0.92** | **0.92** |
| | | Precision | 0.76 | **0.83** | **0.83** | 0.66 | 0.79 | 0.80 |
| | | F1 | 0.75 | **0.87** | 0.77 | 0.73 | 0.85 | 0.85 |
| | Macro | Recall | 0.88 | **0.90** | 0.87 | 0.88 | 0.89 | 0.89 |
| | | Precision | 0.87 | **0.88** | **0.88** | 0.85 | 0.87 | 0.87 |
| | | F1 | 0.87 | **0.88** | 0.87 | 0.86 | 0.87 | 0.87 |
| $\boldsymbol{k_{cat}}$ (424) | Micro | Recall | 0.60 | 0.76 | 0.60 | 0.66 | **0.80** | **0.80** |
| | | Precision | 0.67 | **0.89** | 0.80 | 0.54 | 0.87 | 0.88 |
| | | F1 | 0.63 | 0.82 | 0.69 | 0.59 | 0.83 | **0.84** |
| | Macro | Recall | 0.87 | 0.89 | 0.85 | 0.82 | 0.89 | **0.91** |
| | | Precision | 0.86 | 0.91 | 0.87 | 0.81 | 0.89 | **0.93** |
| | | F1 | 0.85 | 0.90 | 0.85 | 0.80 | 0.89 | **0.91** |
| | | Recall | 0.48 | 0.66 | 0.52 | 0.42 | **0.68** | 0.67 |

| | | | | | | | | |
|---|---|---|---|---|---|---|---|---|
| ***k*~cat~/K~m~** (438) | Micro | Precision | 0.56 | **0.85** | 0.82 | 0.39 | 0.77 | 0.76 |
| | | F1 | 0.52 | **0.74** | 0.63 | 0.41 | 0.72 | 0.71 |
| | Macro | Recall | 0.82 | **0.89** | 0.85 | 0.75 | 0.86 | 0.88 |
| | | Precision | 0.82 | **0.90** | 0.87 | 0.75 | 0.86 | 0.89 |
| | | F1 | 0.81 | **0.89** | 0.86 | 0.75 | 0.86 | 0.88 |
| **$k_{cleave}$** (54) | Micro | Recall | **0.91** | 0.30 | 0.76 | 0.39 | 0.30 | 0.30 |
| | | Precision | **0.42** | 0.25 | 0.37 | 0.08 | 0.19 | 0.26 |
| | | F1 | **0.57** | 0.27 | 0.49 | 0.14 | 0.23 | 0.28 |
| | Macro | Recall | 0.91 | 0.90 | **0.92** | 0.81 | 0.90 | 0.89 |
| | | Precision | **0.91** | 0.90 | **0.91** | 0.80 | 0.89 | 0.89 |
| | | F1 | 0.91 | 0.90 | **0.92** | 0.80 | 0.89 | 0.89 |

**Supplementary Figure 1:** Ablation study of Claude 3.5 and gpt-4o with varying temperature settings on the annotated dataset of protein enzymes.

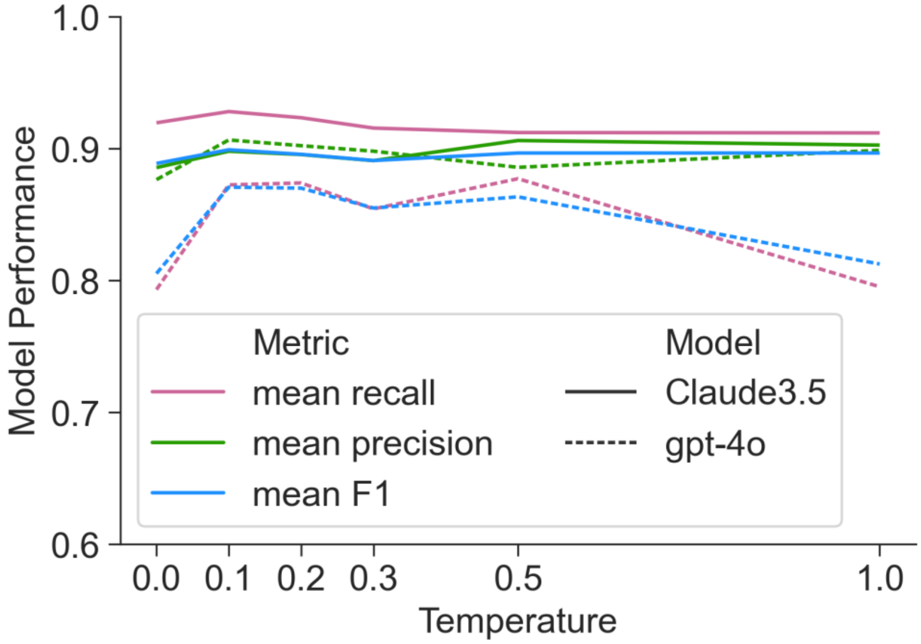

**Supplementary Figure 2:** Open Archive and Open LLM Benchmark for Enzyme Kinetics. **a,** a demo of our pipeline including PDF uploading, OCR, prompt engineering and LLM extraction. **b,** a demo of our annotated dataset which serves as the gold standard for evaluation. **c,** the open archive of enzyme kinetics information. **d-e,** the length distribution of the protein enzyme/ribozyme papers regarding the number of tokens. We use the GPT tokenizer to estimate the number of tokens, but other tokenizers do not make significant differences.

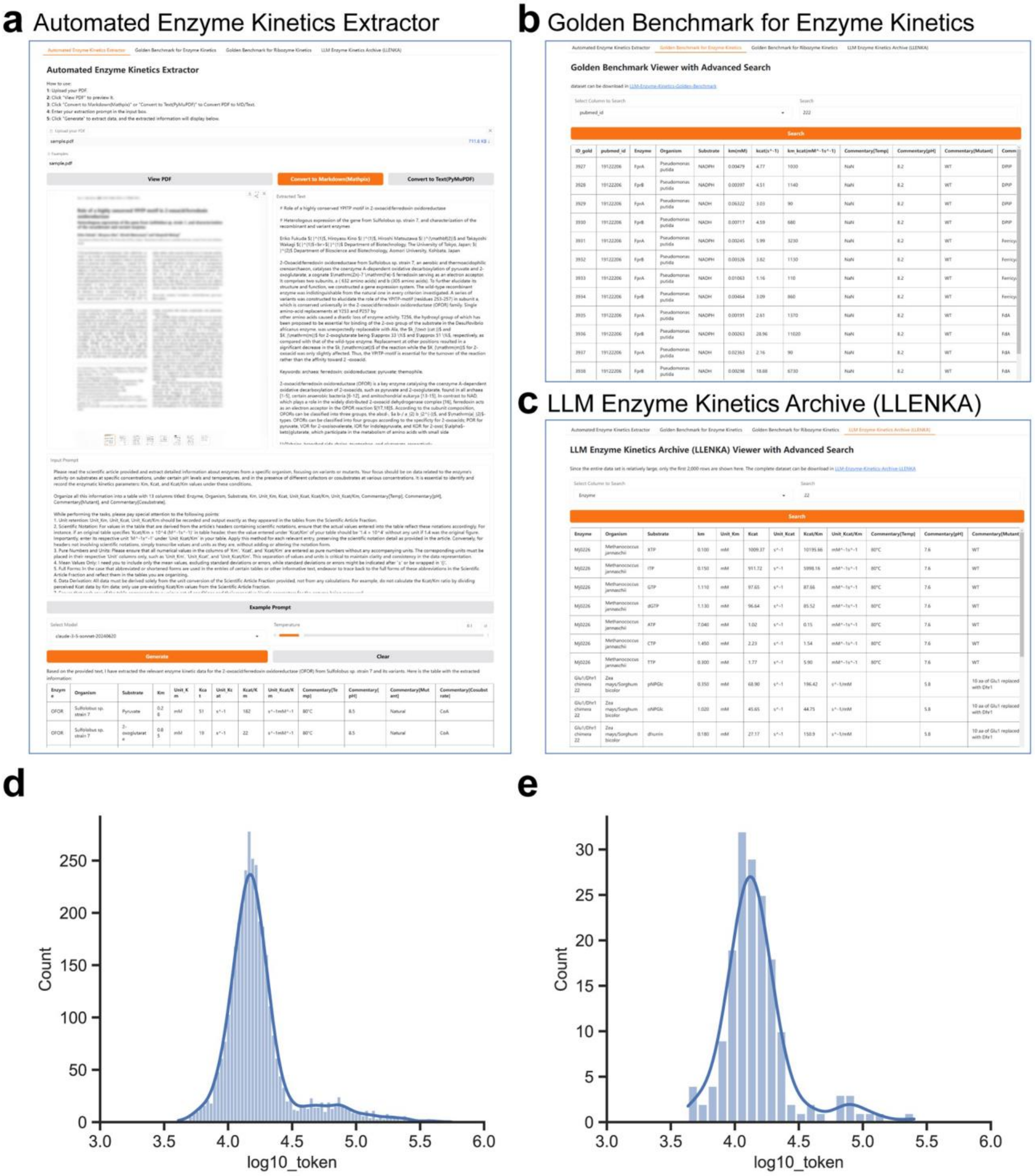

**Supplementary Figure 3:** Ablation study of Claude 3.5 with different OCR software on the annotated dataset of protein enzymes.

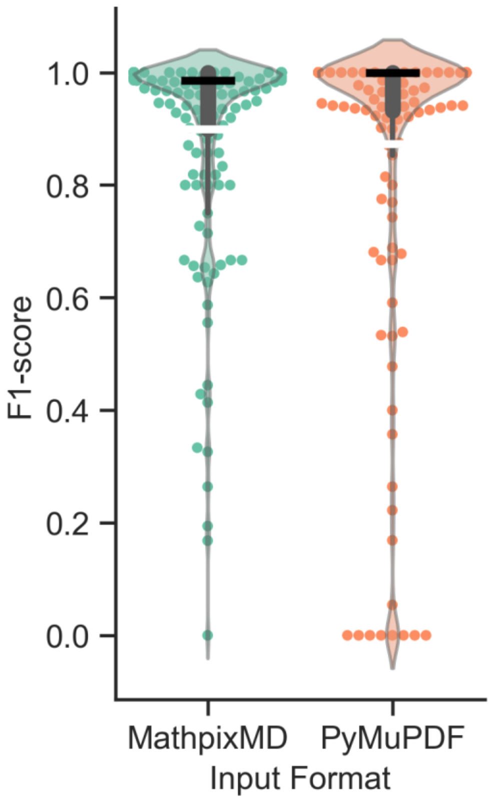

**Supplementary Note 1:** Despite their lower mean and median F1 scores, the other three LLMs (i.e., Qwen, Llama3 and gpt-4o) demonstrate distinct strengths in resolving certain challenging cases encountered by Claude3.5. Here we list three examples.

- Example 1 (PubMed ID: 28411200): Claude3.5 (Precision=0.95, Recall=0.82, F1=0.88) vs Qwen (Precision=1.00, Recall=1.00, F1=1.00)

Claude3.5 extracted results:

| Enzyme | Organism | Substrate | Km | Unit_Km | Kcat | Unit_Kcat | Kcat/Km | Unit_Kcat/Km | Commentary[Temp] | Commentary[pH] | Commentary[Mutant] | Commentary[Cosubstrate] |
|---|---|---|---|---|---|---|---|---|---|---|---|---|
| FNO | Thermobifida fusca | NADH | 14 | mM | 2.2 | s^-1 | 160 | M^-1s^-1 | 25°C | 6.0 | Wild-type | F420 |
| FNO | Thermobifida fusca | NADPH | 7.3 | µM | 3.3 | s^-1 | 450 | mM^-1s^-1 | 25°C | 6.0 | Wild-type | F420 |
| FNO | Thermobifida fusca | NADH | 8.6 | mM | 3.2 | s^-1 | 370 | M^-1s^-1 | 25°C | 6.0 | R51A | F420 |
| FNO | Thermobifida fusca | NADPH | 180 | µM | 1.6 | s^-1 | 6.2 | mM^-1s^-1 | 25°C | 6.0 | R51A | F420 |
| FNO | Thermobifida fusca | NADH | 8.7 | mM | 3.4 | s^-1 | 290 | M^-1s^-1 | 25°C | 6.0 | R51V | F420 |
| FNO | Thermobifida fusca | NADPH | 180 | µM | 1.3 | s^-1 | 9.3 | mM^-1s^-1 | 25°C | 6.0 | R51V | F420 |
| FNO | Thermobifida fusca | NADH | 7.0 | mM | 3.0 | s^-1 | 420 | M^-1s^-1 | 25°C | 6.0 | R55A | F420 |
| FNO | Thermobifida fusca | NADPH | 29 | µM | 8.8 | s^-1 | 300 | mM^-1s^-1 | 25°C | 6.0 | R55A | F420 |
| FNO | Thermobifida fusca | NADH | 6.3 | mM | 2.8 | s^-1 | 440 | M^-1s^-1 | 25°C | 6.0 | R55N | F420 |
| FNO | Thermobifida fusca | NADH | 4.4 | mM | 3.5 | s^-1 | 790 | M^-1s^-1 | 25°C | 6.0 | R55S | F420 |
| FNO | Thermobifida fusca | NADPH | 170 | µM | 6.9 | s^-1 | 41 | mM^-1s^-1 | 25°C | 6.0 | R55S | F420 |
| FNO | Thermobifida fusca | NADH | 9.6 | mM | 3.2 | s^-1 | 330 | M^-1s^-1 | 25°C | 6.0 | R55V | F420 |
| FNO | Thermobifida fusca | NADH | 3.2 | mM | 2.7 | s^-1 | 840 | M^-1s^-1 | 25°C | 6.0 | S50E | F420 |
| FNO | Thermobifida fusca | NADH | 8.2 | mM | 4.2 | s^-1 | 510 | M^-1s^-1 | 25°C | 6.0 | S50Q | F420 |
| FNO | Thermobifida fusca | NADH | 5.0 | mM | 2.6 | s^-1 | 520 | M^-1s^-1 | 25°C | 6.0 | T28A | F420 |
| FNO | Thermobifida fusca | NADPH | 19 | µM | 14 | s^-1 | 720 | mM^-1s^-1 | 25°C | 6.0 | T28A | F420 |
| FNO | Thermobifida fusca | NADH | 10 | mM | 1.6 | s^-1 | 160 | M^-1s^-1 | 25°C | 6.0 | R51ER55A | F420 |
| FNO | Thermobifida fusca | NADH | 6.5 | mM | 2.7 | s^-1 | 420 | M^-1s^-1 | 25°C | 6.0 | R51ER55N | F420 |
| FNO | Thermobifida fusca | NADH | 32 | mM | 4.9 | s^-1 | 150 | M^-1s^-1 | 25°C | 6.0 | R51ER55S | F420 |
| FNO | Thermobifida fusca | NADH | 10 | mM | 2.8 | s^-1 | 280 | M^-1s^-1 | 25°C | 6.0 | R51VR55V | F420 |
| FNO | Thermobifida fusca | NADH | 20 | mM | 2.3 | s^-1 | 120 | M^-1s^-1 | 25°C | 6.0 | S50ER55A | F420 |
| FNO | Thermobifida fusca | NADH | 9.8 | mM | 1.8 | s^-1 | 180 | M^-1s^-1 | 25°C | 6.0 | S50ER55V | F420 |
| FNO | Thermobifida fusca | NADH | 12 | mM | 2.7 | s^-1 | 230 | M^-1s^-1 | 25°C | 6.0 | T28AR51V | F420 |
| FNO | Thermobifida fusca | NADH | 5.4 | mM | 2.5 | s^-1 | 460 | M^-1s^-1 | 25°C | 6.0 | T28AR55A | F420 |
| FNO | Thermobifida fusca | NADPH | 93 | µM | 3.3 | s^-1 | 3.5 | mM^-1s^-1 | 25°C | 6.0 | T28AR55A | F420 |
| FNO | Thermobifida fusca | NADH | 12 | mM | 3.3 | s^-1 | 280 | M^-1s^-1 | 25°C | 6.0 | T28AR51VR55V | F420 |

Qwen extracted results:

| Enzyme | Organism | Substrate | Km | Unit_Km | Kcat | Unit_Kcat | Kcat/Km | Unit_Kcat/Km | Commentary[Temp] | Commentary[pH] | Commentary[Mutant] | Commentary[Cosubstrate] |
|---|---|---|---|---|---|---|---|---|---|---|---|---|
| Tfu-FNO | Thermobifida fusca | NADH | 14 | mM | 2.2 | s^-1 | 160 | M^-1s^-1 | 25°C | 6.0 | WT | None |
| Tfu-FNO | Thermobifida fusca | NADPH | 7.3 | µM | 3.3 | s^-1 | 450 | mM^-1s^-1 | 25°C | 6.0 | WT | None |
| Tfu-FNO | Thermobifida fusca | NADH | 8.6 | mM | 3.2 | s^-1 | 370 | M^-1s^-1 | 25°C | 6.0 | R51A | None |
| Tfu-FNO | Thermobifida fusca | NADPH | >180 | µM | >1.6 | s^-1 | 6.2 | mM^-1s^-1 | 25°C | 6.0 | R51A | None |
| Tfu-FNO | Thermobifida fusca | NADH | 8.7 | mM | 3.4 | s^-1 | 290 | M^-1s^-1 | 25°C | 6.0 | R51V | None |
| Tfu-FNO | Thermobifida fusca | NADPH | >180 | µM | >1.3 | s^-1 | 9.3 | mM^-1s^-1 | 25°C | 6.0 | R51V | None |
| Tfu-FNO | Thermobifida fusca | NADH | 7.0 | mM | 3.0 | s^-1 | 420 | M^-1s^-1 | 25°C | 6.0 | R55A | None |
| Tfu-FNO | Thermobifida fusca | NADPH | 29 | µM | 8.8 | s^-1 | 300 | mM^-1s^-1 | 25°C | 6.0 | R55A | None |
| Tfu-FNO | Thermobifida fusca | NADH | 6.3 | mM | 2.8 | s^-1 | 440 | M^-1s^-1 | 25°C | 6.0 | R55N | None |

```
| Tfu-FNO | Thermobifida fusca | NADPH | >500 | µM | n.d. | s^-1 | n.d. | mM^-1s^-1 | 25°C
| 6.0 | R55N | None |
| Tfu-FNO | Thermobifida fusca | NADH | 4.4 | mM | 3.5 | s^-1 | 790 | M^-1s^-1 | 25°C
| 6.0 | R55S | None |
| Tfu-FNO | Thermobifida fusca | NADPH | 170 | µM | 6.9 | s^-1 | 41 | mM^-1s^-1 | 25°C
| 6.0 | R55S | None |
| Tfu-FNO | Thermobifida fusca | NADH | 9.6 | mM | 3.2 | s^-1 | 330 | M^-1s^-1 | 25°C
| 6.0 | R55V | None |
| Tfu-FNO | Thermobifida fusca | NADPH | 49 | µM | n.d. | s^-1 | n.d. | mM^-1s^-1 | 25°C
| 6.0 | R55V | None |
| Tfu-FNO | Thermobifida fusca | NADH | 3.2 | mM | 2.7 | s^-1 | 840 | M^-1s^-1 | 25°C
| 6.0 | S50E | None |
| Tfu-FNO | Thermobifida fusca | NADPH | >500 | µM | n.d. | s^-1 | n.d. | mM^-1s^-1 | 25°C
| 6.0 | S50E | None |
| Tfu-FNO | Thermobifida fusca | NADH | 8.2 | mM | 4.2 | s^-1 | 510 | M^-1s^-1 | 25°C
| 6.0 | S50Q | None |
| Tfu-FNO | Thermobifida fusca | NADPH | >500 | µM | n.d. | s^-1 | n.d. | mM^-1s^-1 | 25°C
| 6.0 | S50Q | None |
| Tfu-FNO | Thermobifida fusca | NADH | 5.0 | mM | 2.6 | s^-1 | 520 | M^-1s^-1 | 25°C
| 6.0 | T28A | None |
| Tfu-FNO | Thermobifida fusca | NADPH | 19 | µM | 14 | s^-1 | 720 | mM^-1s^-1 | 25°C
| 6.0 | T28A | None |
| Tfu-FNO | Thermobifida fusca | NADH | 10 | mM | 1.6 | s^-1 | 160 | M^-1s^-1 | 25°C
| 6.0 | R51ER55A | None |
| Tfu-FNO | Thermobifida fusca | NADPH | >500 | µM | n.d. | s^-1 | n.d. | mM^-1s^-1 | 25°C
| 6.0 | R51ER55A | None |
| Tfu-FNO | Thermobifida fusca | NADH | 6.5 | mM | 2.7 | s^-1 | 420 | M^-1s^-1 | 25°C
| 6.0 | R51ER55N | None |
| Tfu-FNO | Thermobifida fusca | NADPH | >500 | µM | n.d. | s^-1 | n.d. | mM^-1s^-1 | 25°C
| 6.0 | R51ER55N | None |
| Tfu-FNO | Thermobifida fusca | NADH | 32 | mM | 4.9 | s^-1 | 150 | M^-1s^-1 | 25°C
| 6.0 | R51ER55S | None |
| Tfu-FNO | Thermobifida fusca | NADPH | >500 | µM | n.d. | s^-1 | n.d. | mM^-1s^-1 | 25°C
| 6.0 | R51ER55S | None |
| Tfu-FNO | Thermobifida fusca | NADH | 10 | mM | 2.8 | s^-1 | 280 | M^-1s^-1 | 25°C
| 6.0 | R51VR55V | None |
| Tfu-FNO | Thermobifida fusca | NADPH | >500 | µM | n.d. | s^-1 | n.d. | mM^-1s^-1 | 25°C
| 6.0 | R51VR55V | None |
| Tfu-FNO | Thermobifida fusca | NADH | 20 | mM | 2.3 | s^-1 | 120 | M^-1s^-1 | 25°C
| 6.0 | S50ER55A | None |
| Tfu-FNO | Thermobifida fusca | NADPH | >500 | µM | n.d. | s^-1 | n.d. | mM^-1s^-1 | 25°C
| 6.0 | S50ER55A | None |
| Tfu-FNO | Thermobifida fusca | NADH | 9.8 | mM | 1.8 | s^-1 | 180 | M^-1s^-1 | 25°C
| 6.0 | S50ER55V | None |
| Tfu-FNO | Thermobifida fusca | NADPH | >500 | µM | n.d. | s^-1 | n.d. | mM^-1s^-1 | 25°C
| 6.0 | S50ER55V | None |
| Tfu-FNO | Thermobifida fusca | NADH | 12 | mM | 2.7 | s^-1 | 230 | M^-1s^-1 | 25°C
| 6.0 | T28AR51V | None |
| Tfu-FNO | Thermobifida fusca | NADPH | >500 | µM | n.d. | s^-1 | n.d. | mM^-1s^-1 | 25°C
| 6.0 | T28AR51V | None |
| Tfu-FNO | Thermobifida fusca | NADH | 5.4 | mM | 2.5 | s^-1 | 460 | M^-1s^-1 | 25°C
| 6.0 | T28AR55A | None |
| Tfu-FNO | Thermobifida fusca | NADPH | 93 | µM | 3.3 | s^-1 | 3.5 | mM^-1s^-1 | 25°C
| 6.0 | T28AR55A | None |
| Tfu-FNO | Thermobifida fusca | NADH | 12 | mM | 3.3 | s^-1 | 280 | M^-1s^-1 | 25°C
| 6.0 | T28AR51VR55V | None |
| Tfu-FNO | Thermobifida fusca | NADPH | >500 | µM | n.d. | s^-1 | n.d. | mM^-1s^-1 | 25°C
| 6.0 | T28AR51VR55V | None |
```

- Example 2 (PubMed ID: 26154028): Claude3.5 (Precision=0.58, Recall=0.94, F1=0.71) vs Llama3 (Precision=0.64, Recall=1.00, F1=0.78)

Claude3.5 extracted results:

```
| Enzyme | Organism | Substrate | Km | Unit_Km | Kcat | Unit_Kcat | Kcat/Km | Unit_Kcat/Km | Commentary[Temp] |
Commentary[pH] | Commentary[Mutant] | Commentary[Cosubstrate] |
|---------|----------|-----------|-----|---------|------|-----------|---------|--------------|------------------|------
----------|-------------------|----------------------|
| Homoserine dehydrogenase | Pyrococcus horikoshii | Homoserine | 6.1 | mM | 70.1 | s^-1 | 219 | s^-1·mM^-1 | 50°C | 11.0
| Wild type | NAD |
```

| Homoserine dehydrogenase | Pyrococcus horikoshii | NAD | 0.32 | mM | 70.1 | s^-1 | 219 | s^-1·mM^-1 | 50°C | 11.0 | Wild type | Homoserine |
| Homoserine dehydrogenase | Pyrococcus horikoshii | Homoserine | 0.95 | mM | 96.1 | s^-1 | 102 | s^-1·mM^-1 | 50°C | 11.0 | R40A | NAD |
| Homoserine dehydrogenase | Pyrococcus horikoshii | NAD | 0.95 | mM | 96.1 | s^-1 | 102 | s^-1·mM^-1 | 50°C | 11.0 | R40A | Homoserine |
| Homoserine dehydrogenase | Pyrococcus horikoshii | NADP | 0.04 | mM | 3.4 | s^-1 | 89.1 | s^-1·mM^-1 | 50°C | 11.0 | R40A | Homoserine |
| Homoserine dehydrogenase | Pyrococcus horikoshii | Homoserine | 0.05 | mM | 48.5 | s^-1 | 1020 | s^-1·mM^-1 | 50°C | 11.0 | K57A | NAD |
| Homoserine dehydrogenase | Pyrococcus horikoshii | NAD | 0.05 | mM | 48.5 | s^-1 | 1020 | s^-1·mM^-1 | 50°C | 11.0 | K57A | Homoserine |
| Homoserine dehydrogenase | Pyrococcus horikoshii | NADP | 0.06 | mM | 26.7 | s^-1 | 518 | s^-1·mM^-1 | 50°C | 11.0 | K57A | Homoserine |
| Homoserine dehydrogenase | Pyrococcus horikoshii | Aspartate semialdehyde | 0.019 | mM | 295 | μmol·min^-1·mg^-1 | 15526 | μmol·min^-1·mg^-1·mM^-1 | 50°C | 7.5 | Wild type | NADH |
| Homoserine dehydrogenase | Pyrococcus horikoshii | Aspartate semialdehyde | 0.015 | mM | 8.7 | μmol·min^-1·mg^-1 | 580 | μmol·min^-1·mg^-1·mM^-1 | 50°C | 7.5 | Wild type | NADPH |

Llama3 extracted results:

| Enzyme | Organism | Substrate | Km | Unit_Km | Kcat | Unit_Kcat | Kcat/Km | Unit_Kcat/Km | Commentary[Temp] | Commentary[pH] | Commentary[Mutant] | Commentary[Cosubstrate] |
| --- | --- | --- | --- | --- | --- | --- | --- | --- | --- | --- | --- | --- |
| HseDH | Pyrococcus horikoshii | Hse | 6.1 | mM | 70.1 | s^-1 | 219 | s^-1mM^-1 | 50°C | 11.0 | WT | NAD^+ |
| HseDH | Pyrococcus horikoshii | NAD | 0.32 | mM | 70.1 | s^-1 | 219 | s^-1mM^-1 | 50°C | 11.0 | WT | Hse |
| HseDH | Pyrococcus horikoshii | NADP | - | - | 0 | s^-1 | - | - | 50°C | 11.0 | WT | Hse |
| HseDH | Pyrococcus horikoshii | NAD | 0.95 | mM | 96.1 | s^-1 | 102 | s^-1mM^-1 | 50°C | 11.0 | R40A | Hse |
| HseDH | Pyrococcus horikoshii | NADP | 0.04 | mM | 3.4 | s^-1 | 89.1 | s^-1mM^-1 | 50°C | 11.0 | R40A | Hse |
| HseDH | Pyrococcus horikoshii | NAD | 0.05 | mM | 48.5 | s^-1 | 1020 | s^-1mM^-1 | 50°C | 11.0 | K57A | Hse |
| HseDH | Pyrococcus horikoshii | NADP | 0.06 | mM | 26.7 | s^-1 | 518 | s^-1mM^-1 | 50°C | 11.0 | K57A | Hse |
| HseDH | Pyrococcus horikoshii | Asa | - | - | 295 | s^-1 | - | - | 50°C | 7.5 | WT | NADH |
| HseDH | Pyrococcus horikoshii | Asa | - | - | 8.7 | s^-1 | - | - | 50°C | 7.5 | WT | NADPH |
| HseDH | Pyrococcus horikoshii | NADH | 0.019 | mM | 295 | s^-1 | - | - | 50°C | 7.5 | WT | Asa |
| HseDH | Pyrococcus horikoshii | NADPH | 0.015 | mM | 8.7 | s^-1 | - | - | 50°C | 7.5 | WT | Asa |

- Example 3 (PubMed ID: 17715131): Claude3.5 (Precision=1.00, Recall=0.67, F1=0.80) vs gpt-4o (Precision=1.00, Recall=0.96, F1=0.98)

Claude3.5 extracted results:

| Enzyme | Organism | Substrate | Km | Unit_Km | Kcat | Unit_Kcat | Kcat/Km | Unit_Kcat/Km | Commentary[Temp] | Commentary[pH] | Commentary[Mutant] | Commentary[Cosubstrate] |
| --- | --- | --- | --- | --- | --- | --- | --- | --- | --- | --- | --- | --- |
| EAH | Nicotiana tabacum | EA | 19.2 | μM | 0.5 | s^-1 | 0.03 | | | | | NADPH |
| EAH | Nicotiana tabacum | 1β-(OH)EA | 1.7 | μM | 0.6 | s^-1 | 0.35 | | | | | NADPH |
| HPO | Hyoscyamus muticus | PSD | 14.0 | μM | 2.1 | s^-1 | 0.15 | | | | | NADPH |
| HPO | Hyoscyamus muticus | PSD | 1.7 | μM | 0.1 | s^-1 | 0.06 | | | | | NADPH |
| HPO | Hyoscyamus muticus | Solavetivol | 1.2 | μM | 0.1 | s^-1 | 0.08 | | | | | NADPH |
| HPO | Hyoscyamus muticus | Valencene | 11.5 | μM | 0.1 | s^-1 | 0.01 | | | | | NADPH |
| HPO | Hyoscyamus muticus | Valencene | 7.4 | μM | 1.9 | s^-1 | 0.26 | | | | | NADPH |
| HPO | Hyoscyamus muticus | EA | 3.3 | μM | 0.2 | s^-1 | 0.06 | | | | | NADPH |
| HPO | Hyoscyamus muticus | EE | 7.8 | μM | 0.8 | s^-1 | 0.10 | | | | | NADPH |
| HPO | Hyoscyamus muticus | Cedr-8-ene | 26.5 | μM | 0.1 | s^-1 | 0.00 | | | | | NADPH |
| EAH | Nicotiana tabacum | PSD | 9.2 | μM | 0.4 | s^-1 | 0.04 | | | | | NADPH |
| EAH | Nicotiana tabacum | PSD | 11.4 | μM | 0.6 | s^-1 | 0.05 | | | | S368V | NADPH |
| EAH | Nicotiana tabacum | PSD | 7.6 | μM | 0.4 | s^-1 | 0.05 | | | | S368V/I484V | NADPH |

| | | | | | | | | | | | |
|---|---|---|---|---|---|---|---|---|---|---|---|
| EAH | Nicotiana tabacum | PSD | 6.5 | µM | 0.6 | s^-1 | 0.09 | | | S368V/S482V | NADPH |
| HPO | Hyoscyamus muticus | PSD | 6.2 | µM | 3.8 | s^-1 | 0.61 | | | V482I | NADPH |
| HPO | Hyoscyamus muticus | Valencene | 6.9 | µM | 5.1 | s^-1 | 0.74 | | | V482I | NADPH |
| HPO | Hyoscyamus muticus | EA | 17.9 | µM | 1.4 | s^-1 | 0.08 | | | V482I | NADPH |
| HPO | Hyoscyamus muticus | PSD | 8.4 | µM | 12.7 | s^-1 | 1.51 | | | V480I/A484I | NADPH |
| HPO | Hyoscyamus muticus | Valencene | 19.2 | µM | 5.7 | s^-1 | 0.30 | | | V480I/A484I | NADPH |
| HPO | Hyoscyamus muticus | EA | 2.5 | µM | 0.7 | s^-1 | 0.28 | | | V480I/A484I | NADPH |
| HPO | Hyoscyamus muticus | PSD | 13.0 | µM | 20.7 | s^-1 | 1.59 | | | V482I/A484I | NADPH |
| HPO | Hyoscyamus muticus | Valencene | 12.1 | µM | 15.9 | s^-1 | 1.31 | | | V482I/A484I | NADPH |
| HPO | Hyoscyamus muticus | EA | 4.3 | µM | 2.8 | s^-1 | 0.65 | | | V482I/A484I | NADPH |

### gpt-4o extracted results:

| Enzyme | Organism | Substrate | Km | Unit_Km | Kcat | Unit_Kcat | Kcat/Km | Unit_Kcat/Km | Commentary[Temp] | Commentary[pH] | Commentary[Mutant] | Commentary[Cosubstrate] |
|---|---|---|---|---|---|---|---|---|---|---|---|---|
| EAH | Nicotiana tabacum | 5-epi-aristolochene | 19.2 | µM | 0.5 | s^-1 | 0.03 | µM^-1s^-1 | 30°C | 7.5 | WT | NADPH |
| EAH | Nicotiana tabacum | 1β-hydroxy-5-epi-aristolochene | 1.7 | µM | 0.6 | s^-1 | 0.35 | µM^-1s^-1 | 30°C | 7.5 | WT | NADPH |
| HPO | Hyoscyamus muticus | Premnaspirodiene | 14.0 | µM | 2.1 | s^-1 | 0.15 | µM^-1s^-1 | 30°C | 7.5 | WT | NADPH |
| HPO | Hyoscyamus muticus | Premnaspirodiene | 1.7 | µM | 0.1 | s^-1 | 0.06 | µM^-1s^-1 | 30°C | 7.5 | WT | NADPH |
| HPO | Hyoscyamus muticus | Solavetivol | 1.2 | µM | 0.1 | s^-1 | 0.08 | µM^-1s^-1 | 30°C | 7.5 | WT | NADPH |
| HPO | Hyoscyamus muticus | Valencene | 11.5 | µM | 0.1 | s^-1 | 0.01 | µM^-1s^-1 | 30°C | 7.5 | WT | NADPH |
| HPO | Hyoscyamus muticus | Valencene | 7.4 | µM | 1.9 | s^-1 | 0.26 | µM^-1s^-1 | 30°C | 7.5 | WT | NADPH |
| HPO | Hyoscyamus muticus | 5-epi-aristolochene | 3.3 | µM | 0.2 | s^-1 | 0.06 | µM^-1s^-1 | 30°C | 7.5 | WT | NADPH |
| HPO | Hyoscyamus muticus | 4-epi-eremophilene | 7.8 | µM | 0.8 | s^-1 | 0.10 | µM^-1s^-1 | 30°C | 7.5 | WT | NADPH |
| HPO | Hyoscyamus muticus | Cedr-8-ene | 26.5 | µM | 0.1 | s^-1 | 0.00 | µM^-1s^-1 | 30°C | 7.5 | WT | NADPH |
| EAH | Nicotiana tabacum | Premnaspirodiene | 9.2 | µM | 0.4 | s^-1 | 0.04 | µM^-1s^-1 | 30°C | 7.5 | WT | NADPH |
| EAH, S368V | Nicotiana tabacum | Premnaspirodiene | 11.4 | µM | 0.6 | s^-1 | 0.05 | µM^-1s^-1 | 30°C | 7.5 | S368V | NADPH |
| EAH, S368V/I484V | Nicotiana tabacum | Premnaspirodiene | 7.6 | µM | 0.4 | s^-1 | 0.05 | µM^-1s^-1 | 30°C | 7.5 | S368V/I484V | NADPH |
| EAH, S368V/S482V | Nicotiana tabacum | Premnaspirodiene | 6.5 | µM | 0.6 | s^-1 | 0.09 | µM^-1s^-1 | 30°C | 7.5 | S368V/S482V | NADPH |
| HPO, V482I | Hyoscyamus muticus | Premnaspirodiene | 6.2 | µM | 3.8 | s^-1 | 0.61 | µM^-1s^-1 | 30°C | 7.5 | V482I | NADPH |
| HPO, V480I/A484I | Hyoscyamus muticus | Premnaspirodiene | 8.4 | µM | 12.7 | s^-1 | 1.51 | µM^-1s^-1 | 30°C | 7.5 | V480I/A484I | NADPH |
| HPO, V482I/A484I | Hyoscyamus muticus | Premnaspirodiene | 13.0 | µM | 20.7 | s^-1 | 1.59 | µM^-1s^-1 | 30°C | 7.5 | V482I/A484I | NADPH |
| HPO, V482I | Hyoscyamus muticus | Valencene | 6.9 | µM | 5.1 | s^-1 | 0.74 | µM^-1s^-1 | 30°C | 7.5 | V482I | NADPH |
| HPO, V480I/A484I | Hyoscyamus muticus | Valencene | 19.2 | µM | 5.7 | s^-1 | 0.30 | µM^-1s^-1 | 30°C | 7.5 | V480I/A484I | NADPH |
| HPO, V482I/A484I | Hyoscyamus muticus | Valencene | 12.1 | µM | 15.9 | s^-1 | 1.31 | µM^-1s^-1 | 30°C | 7.5 | V482I/A484I | NADPH |
| HPO, V482I | Hyoscyamus muticus | 5-epi-aristolochene | 17.9 | µM | 1.4 | s^-1 | 0.08 | µM^-1s^-1 | 30°C | 7.5 | V482I | NADPH |
| HPO, V480I/A484I | Hyoscyamus muticus | 5-epi-aristolochene | 2.5 | µM | 0.7 | s^-1 | 0.28 | µM^-1s^-1 | 30°C | 7.5 | V480I/A484I | NADPH |
| HPO, V482I/A484I | Hyoscyamus muticus | 5-epi-aristolochene | 4.3 | µM | 2.8 | s^-1 | 0.65 | µM^-1s^-1 | 30°C | 7.5 | V482I/A484I | NADPH |

**Supplementary Note 2:** We scale our workflow to the full dataset of 3,435 papers from BRENDA where we lack expert-annotated ground truth for performance evaluation. To address this, we propose using BRENDA-recorded entries as an approximate ground truth and comparing the LLM-extracted results against these entries. This approach allows us to extend our analysis from the annotated subset (156 papers) to the full dataset (3,435 papers) and compare the paper-wise F1 distributions (Supplementary Fig. 1). By conducting a Kolmogorov-Smirnov test (see Methods), we could not reject the null hypothesis, indicating that there is insufficient evidence to conclude that the two F1 distributions are different ($p\text{-value} > 0.7$). This statistical result suggests that the performance metrics from the annotated subset are indicative of performance across the larger dataset.

**Supplementary Figure 4:** Performance of Claude 3.5 extracted results evaluated on BRENDA which serves as the approximate ground truth. Left: Annotated dataset (156 papers). Right: Full dataset (3,435 papers).

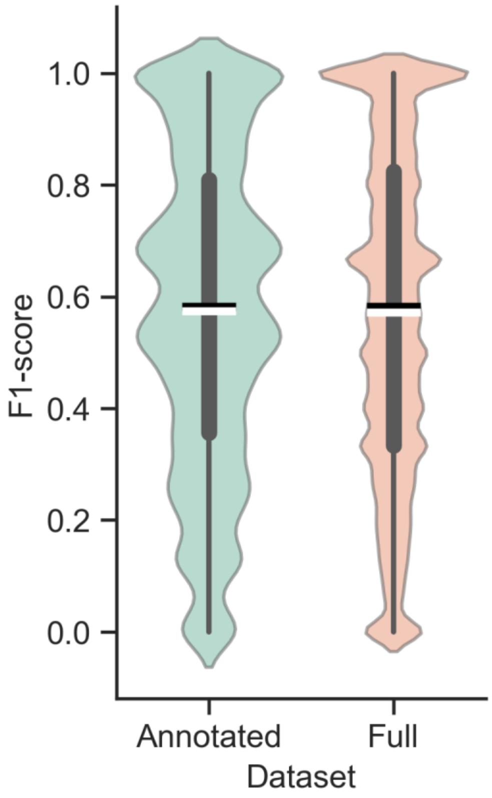

**Supplementary Note 3:** The full version of engineered prompt for a single LLM is as follows:

"Please read the scientific article provided and extract detailed information about enzymes from a specific organism, focusing on variants or mutants. Your focus should be on data related to the enzyme's kinetics on substrates at specific concentrations, under certain pH levels and temperatures, and in the presence of different cofactors or cosubstrates at various concentrations. It is essential to identify and record the enzymatic kinetics parameters: Km, Kcat, and Kcat/Km values under these conditions.

Organize all this information into a table with 13 columns titled: Enzyme, Organism, Substrate, Km, Unit_Km, Kcat, Unit_Kcat, Kcat/Km, Unit_Kcat/Km, Commentary[Temp], Commentary[pH], Commentary[Mutant], and Commentary[Cosubstrate].

While performing the tasks, please pay special attention to the following points:

1. Unit retention: Unit_Km, Unit_Kcat, Unit_Kcat/Km should be recorded and output exactly as they appeared in the tables from the Scientific Article Fraction.
2. Scientific Notation: For values in the table that are derived from the article's headers containing scientific notations, ensure that the actual values entered into the table reflect these notations accordingly. For instance, if an original table specifies 'Kcat/Km × 10^4 (M^-1s^-1)' in table header, then the value entered under 'Kcat/Km' of your table should be '1.4 × 10^4' without any unit if 1.4 was the original figure. Importantly, enter its respective unit 'M^-1s^-1' under 'Unit_Kcat/Km' in your table. Apply this method for each relevant entry, preserving the scientific notation detail as provided in the article. Conversely, for headers not involving scientific notations, simply transcribe values and units as they are, without adding or altering the notation form.
3. Pure Numbers and Units: Please ensure that all numerical values in the columns of 'Km', 'Kcat', and 'Kcat/Km' are entered as pure numbers without any accompanying units. The corresponding units must be placed in their respective 'Unit' columns only, such as 'Unit_Km', 'Unit_Kcat', and 'Unit_Kcat/Km'. This separation of values and units is critical to maintain clarity and consistency in the data representation.
4. Mean Values Only: I need you to include only the mean values, excluding standard deviations

or errors, while standard deviations or errors might be indicated after '±' or be wrapped in '()'.

5. Full Forms: In the case that abbreviated or shortened forms are used in the entries of certain tables or other informative text, endeavor to trace back to the full forms of these abbreviations in the Scientific Article Fraction and reflect them in the tables you are organizing.
6. Data Derivation: All data must be derived solely from the unit conversion of the Scientific Article Fraction provided, not from any calculations. For example, do not calculate the Kcat/Km ratio by dividing perceived Kcat data by Km data; only use pre-existing Kcat/Km values from the Scientific Article Fraction.
7. Ensure that each row of the table corresponds to a unique set of conditions and their respective kinetic parameters for the enzyme being measured.

Output the table using the pipe symbol (|) as the delimiter, ensuring each entry is separated by a pipe symbol and properly aligned to maintain the structure of the table. I need you to include only the mean values, excluding standard deviations or errors, while standard deviations or errors might be indicated after '±' or be wrapped in '()'. Include all details and rows in the output, providing a comprehensive extraction of every data point without omissions. Format the complete table data clearly, ensuring that every piece of information is included and no data points are left out. Do not use ellipses or any other form of indication suggesting information is continued elsewhere. The full dataset must be provided as per the structure above, ensuring the integrity and usability of the data for subsequent analyses or applications. Present the complete table data in a clear and organized format in your response, without the need for further confirmation or prompts.

Please pay attention to the pipe format as shown in the example below. This format is for reference only regarding the structure; the content within is not the focus of this instruction.

| Enzyme | Organism | Substrate | Km | Unit_Km | Kcat | Unit_Kcat | Kcat/Km | Unit_Kcat/Km | Commentary[Temp] | Commentary[pH] | Commentary[Mutant] | Commentary[Cosubstrate] |
| --- | --- | --- | --- | --- | --- | --- | --- | --- | --- | --- | --- | --- |

| | | | | | | | | | | | | |
|---|---|---|---|---|---|---|---|---|---|---|---|---|
| Enzyme1 | Bacillus subtilis | Substrate_A | 7.3 | mM | 6.4 | s^-1 | 1.4 × 10^4 | M^-1s^-1 | 37°C | 5.0 | WT | NADP^+ |
| Enzyme2 | Escherichia coli | Substrate_B | 5.9 | mM | 9.8 | s^-1 | 29000 | mM^-1min^-1 | 60°C | 10.0 | Q176E | NADPH |
| Enzyme3 | Homo sapiens | Substrate_C | 6.9 | mM | 15.6 | s^-1 | 43000 | µM^-1s^-1 | 65°C | 8.0 | T253S | NAD^+ |

Scientific Article Fraction :

'''

{{Scientific Article Fraction}}

'''

Structure your responses to allow for seamless concatenation, presenting all tabular data from a scientific article as a single table, even if the original content had multiple tables. Use the full response capacity to maximize data presentation, avoiding summarizations, commentaries, or introductions at the end of each response. The subsequent response should pick up precisely where the preceding one concluded, commencing from the following character, without the necessity to reiterate the table header or the fragmented words. This method ensures the table is presented completely and seamlessly, despite character limit constraints. I will prompt with 'continue' for continuation when necessary. Please start by outputting the first segment of the table according to these guidelines."

**Supplementary Note 4:** The full version of prompt of the aggregation agent is as follows, here [LLM extraction prompt] refers to the prompt introduced in **Supplementary Note 3**:

"The following is a [scientific article], please read it carefully: {provided_file_content}.

And the corresponding [LLM extraction prompt]: {full version of engineered prompt for a single LLM}.

Next are the responses of the four LLMs:

[response_gpt-4o]: {gpt_4o_response}.

[response_claude-3-5-sonnet-20240620]: {claude_response}.

[response_Meta-Llama-3.1-405B-Instruct]: {llama_response}.

[response_qwen-plus-0806]: {qwen_response}.

Please check these [responses of the four LLMs] according to the provided [scientific article], [LLM extraction prompt] and organize them into a final table."
